# Multi-band component analysis for EEG artifact removal and source reconstruction with application to gamma-band activity

**DOI:** 10.1101/205120

**Authors:** Yaqub Jonmohamadi, Suresh D. Muthukumaraswamy

## Abstract

Independent component analysis (ICA) has been used extensively for artifact removal and reconstruction of neuronal time-courses in electroencephalography (EEG). Typically, ICA is applied on wide-band EEG (for example 1 to 100 Hz or similar ranges). Since EEG captures the activities of a large number of sources and the fact that number of the components separated by the ICA is limited by the number of the sensors, only the stronger sources (in terms of magnitude and duration) will be detected by the ICA, and the activity of weaker sources will be lost or scattered amongst the stronger components. Because of the 1/f nature of the EEG spectra this biases components to the lower frequency ranges. Here we used a versatile combination of a filter bank, PCA and ICA, calling it multi-band ICA, to both increase the number of the ICA components substantially and improve the SNR of the separated components. Using band-pass filtering we break the original signal mixture into several subbands, and using PCA we reduce the dimensionality of each subband, before applying ICA to a matrix containing all the principal components from each band. Using simulated sources and real EEG of participants, we demonstrate that multi-band ICA is able to outperform the traditional wide-band ICA in terms of both signal-noise ratio of the separated sources and the number of the identified independent components. We successfully separated the gamma-band neuronal components time-locked to a visual stimulus, as well as weak sources which are not detectable by wide-band ICA.

## 1. Introduction

Electroencephalography (EEG) is the scalp recording of neuroelectrical activities. Due to its millisecond temporal resolution, simplicity, and low cost, it is the most popular technique for the study of brain dynamics. These advantages come with a cost, that is the EEG signals are contaminated with various types of noise and artifacts, that makes physiological interpretation of raw EEG extremely difficult. Hence, advanced signal processing techniques are necessary in order to extract meaningful information from the recordings. Typically, the EEG is divided into several frequency bands, delta (0-4 Hz), theta (4-8 Hz), alpha (8-12 Hz), beta (12-30 Hz), and gamma (30 to 100 Hz). The most common known artifacts are ocular and cardiac artifacts, muscle, sensor, and line noise. Each of these has a certain band-width, for example, ocular artifacts (eye blinks and movement) are predominantly in the lower frequencies while the muscle artifact is scattered in the beta and gamma bands. Numerous techniques have been been proposed to remove these artifacts form the EEG, for example, using regression of the electrooculographic (EOG) signal to remove the ocular artifact [17, 50, 14, 16, 51]. However, there are a few deficiencies associated with the regressors, making their application suboptimal, for example, the simple regressors may overcompensate the ocular artifact and produce new artifacts due to the differences between the EOG and EEG transfer function [48, 41, 28]. Another problem is the cancellation of the common neuronal information between the EOG and EEG [28]. Moreover, the application of the regressors is limited by noise which can be picked up by a reference electrode [44], hence not every artifact can be eliminated by regression, for example, muscle artifact. Nevertheless, the adaptive methods [1, 38] are still the most effective tools for rejection of certain artifacts such as gradient artifacts which is induced by the magnetic resonance imaging (MRI) machine during concurrent EEG/fMRI (functional MRI) recording.

Spatiotemporal filtering using a dipole model was proposed by Berg and Scherg [5] as an approach for removal of the ocular artifact. Its major limitation is the reliance on a prior specification of the number and location of the dipoles [31]. As a more effective alternative to dipole modeling, Berg and Scherg [6] proposed use of principal component analysis (PCA) for eye artifact removal. However, components separated by the PCA show the directions with maximum variance.

This means weak neural sources could be scattered among strong artifacts separated by the PCA, and vice versa, that is, weak noises could exist among the principal components representing strong neural sources, such as, alpha and theta band sources and degrade signal to noise ratio (SNR) of such components. Hence, the removal of PCA components inevitably results in removal of the neural sources as illustrated by Lagerlund et al. [30]. Unlike PCA, independent component analysis (ICA) maximises the non-gaussianity of the components by measures such as entropy or kurtosis. While there are several algorithms proposed for ICA, generally speaking, ICA is based on a few assumptions, most importantly: (1) sources are linearly mixed, (2) sources are independent from each other, and (3) the number of the sources is equivalent to the number of the signal mixtures. There have been variants of ICA which focus on relaxing one of the assumptions in order to optimise the ICA for a certain application. Here, our approach is concerned with the number of the sources relative to the number of the signal mixtures, that is, the number of the sources could be smaller or greater than the number of the signal mixtures. This uncertainty regarding the true number of the sources in relation to the number of the signal mixtures has been addressed in case of EEG [34].

Since ICA was first applied for EEG [34], it has become the most extensively used technique, both in EEG and magnetoencephalography (MEG), for removal of artifacts and reconstruction of neural time-courses both in sensor space [39, 28, 33, 22, 46, 29, 27, 28, 13], and recently reconstruction of the neural source in the source space [7, 32, 2, 25, 24, 26, 18]. Although ICA separates sources using statistical properties of the data, we will show that the performance of the ICA can be affected by the relative power distribution of the EEG sources, that is, activities in delta and alpha bands are several times stronger than the sources in the gamma band. It is also highly likely that the number of the actual sources recorded by EEG or MEG is greater than the number of the signal mixtures. Moreover, the number of ICA components is upper bounded by the number of the signal mixtures, meaning only the strongest sources will dominate the ICA outputs and the weaker sources will be scattered among them. This means application of the ICA over the wide-band EEG and rejection of the artifact/noise components could result in loss of neuroelectrical sources. For example the high frequency neural sources in gamma-band are weak and could be scattered among the stronger sources separated by wide-band ICA. Furthermore, there are strong sources of noise in the gamma-band such as facial and neck muscles which makes it a challenging situation for the study of gamma-band neural sources [15, 49, 52, 19, 36, 37].

Subband ICA [45, 35, 11] is an alternative to the standard (wide-band) ICA in which the signal mixtures are band-pass filtered using a filter bank, then the ICA is applied on each band separately and the independence of each band is measured using a proposed criteria such as “performance index” or kurtosis. The filter bank separates the sources spectrally and reduces the burden of the decomposition from the ICA. Here we introduce multi-band ICA as an alternative to the standard wide-band ICA which is similar to the subband ICA in terms of applying the bandpass filtering before signal decomposition, but is different in terms of applying the ICA, that is rather than applying the ICA to each band, we apply PCA to each band and only pass a defined number of principal components of each band to produce a global matrix of principal components (PCs) which represents the PCs of all the bands. In this way, the PCA finds the maximum variance of each band separately, hence the sources of one band, for example, gamma, are not compared with the sources from another band such as delta, which is the case in wide-band ICA. Then we apply ICA to reconstruct the independent components (ICs). The advantage of this approach over the subband ICA is that we are not introducing any new criteria such as “performance index” to decide which bands are independent and let the ICA decides on remixing the principal components of different bands. We note this remixing between the subbands via ICA depends on which flavor of ICA is used and will be discussed in later sections. In comparison to the wide-band ICA, the multi-band ICA can reconstruct weak sources which have little chance to be reconstructed via wide-band ICA and instead would be scattered among the stronger ICs. We also demonstrate that sources reconstructed by mutiband ICA have a higher SNR compared with the wide-band ICA. Similar to [12] who showed that it is possible to extract more than one source from a single channel EEG, we will show that the number of the extracted sources is not upper-bounded by the number of the EEG channels. The novelty of this approach is in terms of order of the application of popular signal decomposition techniques such as ICA, PCA and filter banks, rather than introducing new selection and measurement criteria. This offers ease of application and we have provided the source code [23] for multi-band ICA with sample EEG and simulated sources, implemented in MATLAB, using the FieldTrip toolbox and data structure [40].

In the next two sections, we describe the theoretical background, simulations and setups for the acquisition of real EEG data. Throughout this paper, plain italics refers scalars, lower-case boldface italics refers vectors, and upper-case boldface italics indicate matrices.

## 2. Methods

### 2.1. Band pass filtering

The recorded MEG or EEG signal for *K* time samples on *M* sensors, ***X*** ∈ ℜ^*M×K*^, can be written as

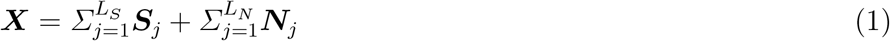

where the *L*_*S*_ and *L*_*N*_ refer to the number of the brain and artifact/noise sources, respectively. The true number of the sources is not known [34] and it depends on several factors, including the duration of the recordings, individual differences between participants, and the task that the subject is performing. Hence, the total number of sources could be more or less than the number of the sensors. The use of wide-band ICA limits the number of the detected ICs by the number of channels (*M*). This is one of the disadvantages of the wide-band ICA over the multi-band ICA. Using a filter bank we can split the wide-band EEG or MEG into several subbands,

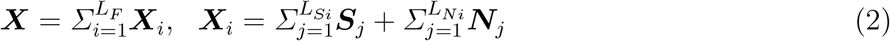

where the *L*_*F*_ refers to the number of the subband (typically delta, theta, alpha, beta, lower gamma, higher gamma, i.e., *L*_*F*_ = 6). The *L*_*Si*_ and *L*_*Ni*_ refer to the number of the brain and noise sources in the *ith* band.

### 2.2. PCA

Since, we do not know what *L*_*Si*_ and *L*_*Ni*_ are, and at the same time there is a strong possibility that each subband contains a smaller number of combined sources than the number of the sensors, i.e., *M* ≥ *L*_*Si*_ + *L*_*Ni*_, particularly when there are large number of sensors, for example 64 or more, it is necessary to reduce the size of the time-courses in each subband to avoid the undercomplete ICA problem. Using singular value decomposition (SVD) to break the signals into PCs, we can reduce the size of the data matrix in each subband to a subsapce containing the first few PCs of that subband:

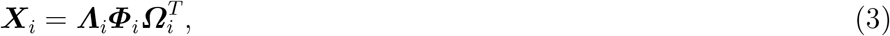

where ***Λ***_*i*_ is an *M* × *M* matrix containing the normalised topographic maps of principal components, and 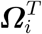 is a *K* × *K* matrix containing the normalised time-courses of the principal components, and ***Φ***_*i*_ is an *M* × *K* diagonal matrix with the diagonal elements being the singular values in descending order. Here, we need to keep the first few PCs of ***Λ***_*i*_ and 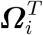. Choosing a small number of components could lead to loss of useful information and a great number may result in an undercomplete ICA problem. There are a number of methods in the literature for choosing an optimal number of components. The most widely use method is the Guttman-Kaiser rule [21], according to which the PCs associated with eigenvalues derived from the data covariance matrix, are kept only if the corresponding eigenvalues are larger in magnitude than the mean of the eigenvalues [8]. However, as the Guttman-Kaiser rule is based on the mean value of the eigenvalues, existence of a large artifact increases the mean of the eigenvalues and can result in substantial change in size of the selected subspace. In noisy EEG applications, the minimum description length (MDL) [42] has been shown shown to be the optimal approach in subspace extraction [47, 9, 10]. Generally, the automatic subspace selection is an extensive topic and depends on several factors beyond the scope this paper. Here we acknowledge using a number between 10 to 15 for subspace extraction in each subband will result in an overall superior performance of the multi-band ICA over the wide-band ICA in terms of reconstruction of weak sources, hence recovering the weak sources. The effects of changing the size of the subspace is discussed in the Results section. Assuming that *N*_*Pi*_ (e.g., *N*_*P*1_ = 10, *N*_*P*2_ = 12,…) is the number of PCs from each band to be extracted, the subspace of each band after dimensional reduction can be written as

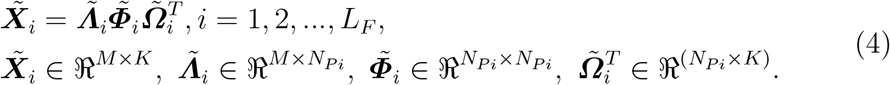

Since we are performing temporal rather than spatial decomposition, we need to create the global matrix of temporal PCs. In order to retain the magnitude order of the temporal PCs, normalised PCs are multiplied with their corresponding singular values

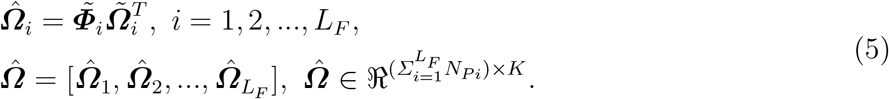

The corresponding global matrix of spatial PCs is

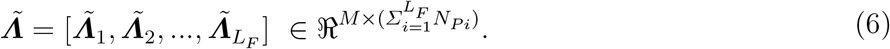

### 2.3. ICA

ICA decomposes the global matrix of the temporal PCs into ICs

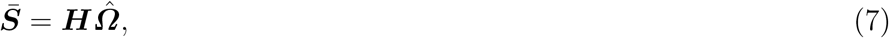

where 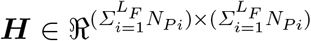 is the unmixing matrix and, ***S̄*** is the matrix of ICs. The topographic maps of 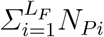 identified components in ***S̄*** can be obtained by multiplication of the global spatial PCs by the mixing matrix

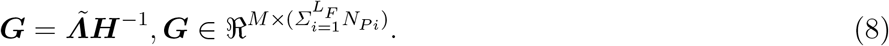

As is shown in equation 8, the number of extracted components is 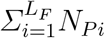 and could be greater or smaller than the number of the channels, for example, if our filter bank has 6 subbands (*L*_*F*_ = 6) and we chose to extract first 15 PCs of each subband (*N*_*P*_ = 15), the total number for the ICs will be 90. The block diagram of the steps in order to perform multi-band ICA is shown in Figure 1.

## 3. Performance evaluation

### 3.1. Simulations

The aim of the simulations is to demonstrate the superiority of the multi-band ICA over the conventional wide-band ICA in terms of the separation power and the SNR of the ICs. We simulated 3 sources and superimposed them on the EEG of a resting state subject. For the simulation of the sources, the boundary element model was used for the computation of the leadfield (template model provided by FieldTrip toolbox). The specifications of the sources together with their topographic maps are represented in Table 1. The resting state EEG was recorded with the sampling rate of 1000 Hz, and after band-passed filtering (0.5-100 Hz) it was downsampled to 500 Hz.

**Figure 1:**
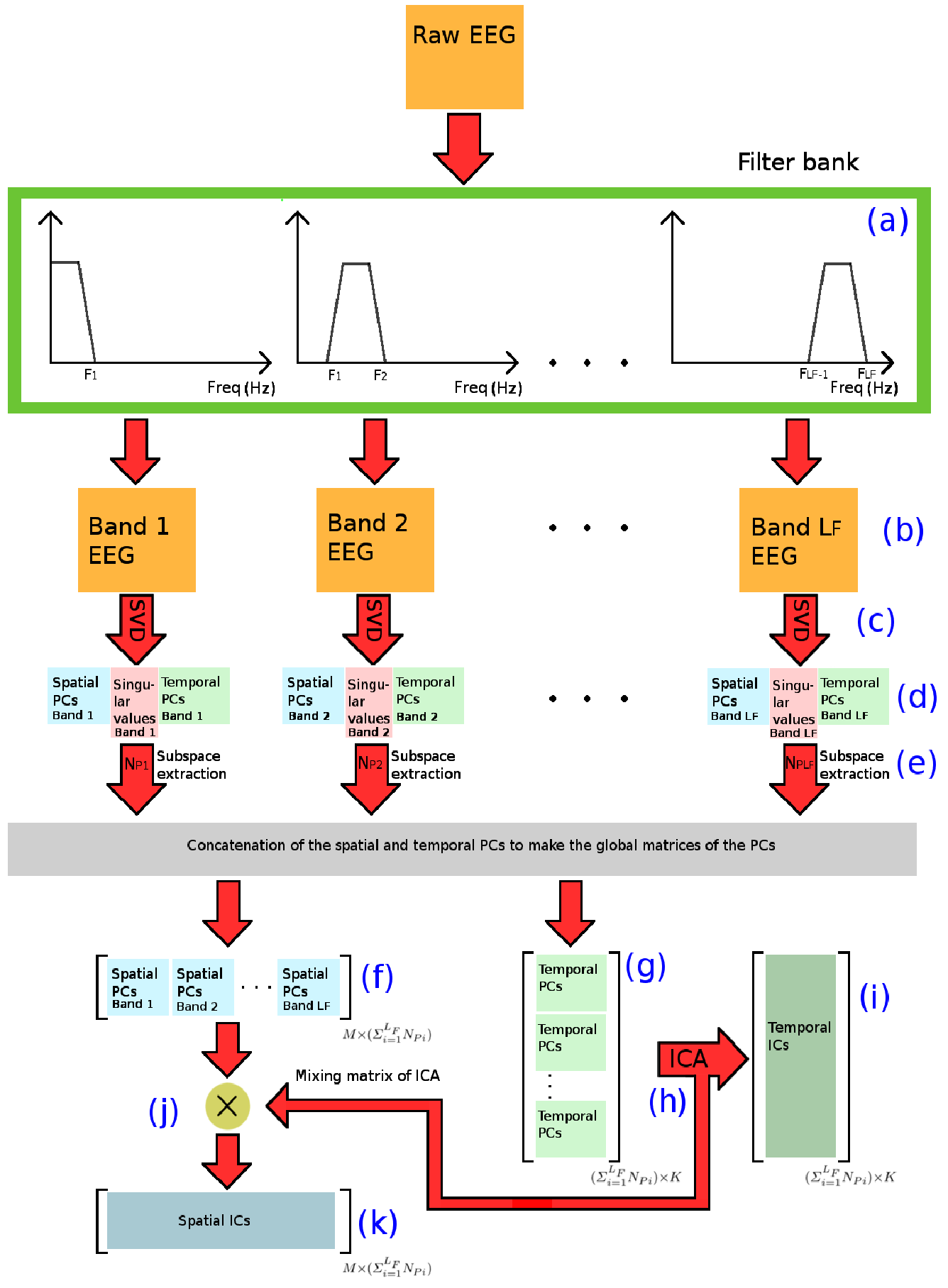
The block diagram of the multi-band ICA. The raw EEG is band passed filtered using the filter banks (a) to create the subband EEGs (b). Using the SVD (c) the subband EEGs are broken into the spatial and temporal subspaces (d) and from each subspace, the first *N*_*Pi*_ is selected (e) to create the global matrices of spatial (f) and temporal (g) principal components. Temporal ICA (h) is then applied to extract the temporal ICs (i), and the mixing matrix of ICA is multiplied (j) by the global matrix of the spatial PCs to create the spatial ICs (k) corresponding to the temporal ICs.

**Table 1:** Details of the simulated sources.

| Source | MNI positions | Moment | Freq. | SNR |
| --- | --- | --- | --- | --- |
| 1 | [0 3 0] cm | [0 1 0] | 45 Hz | 0.04 |
| 2 | [-4 2 2] cm | [0 0 1] | 12 Hz | 0.04 |
| 3 | [0 -6 3] cm | [1 1 1] | 5 Hz | 0.10 |

### 3.2. Reconstruction of visually induced gamma components

EEG data from 22 participants was available. In this study the participants were performing a standard task to induce gamma-band activity in the visual cortex. The stimuli for this task was a black and white annular grating with a spatial frequency of 3 cycles per degree, subtending 16 degrees of visual angle. The grating was presented at 90% contrast, in the centre of the screen, on a grey background. A red dot provided a central fixation point. Participants were seated 90cm from the screen. In this task were two conditions. the stimuli moved inwardly at a rate of 1.33 degrees of visual angle per second. The stimulus was on for between 1-1.1s (pseudorandomly jittered). Participants were instructed to press the spacebar as soon as the stimulus disappeared from the screen. Following a participants response there was then a 1s inter-trial interval. Stimuli were displayed on an ASUS VG248QE computer monitor with a screen resolution of 1920 × 1800 and 144Hz refresh rate generated using the Psychophysics Toolbox. Continuous EEG was recorded using 64 channel Acticap Ag/AgCl active shielded electrodes and Brain Products MRPlus amplifiers. Data were recorded in Brain Vision Recorder (Brain Products GmbH, Germany) with a 1000Hz sampling rate, and 0.1V resolution. After band-pass filtering (0.5-100 Hz) they were downsampled to 500 Hz. FCz was used as an online reference, AFz as ground. Electrode impedance below 10k was achieved prior to recording. Data were first epoched into trials −0.5 ms pre stimulus and 1.5 ms post-stimulus onset. The data were then baselined. Semi-automated artefact rejection was completed using the Fieldtrip toolbox.

Here the aim is to find gamma ICs time-locked to the stimulus. Since the neural activities in the gamma-band are weak, the separation of the gamma-band ICs is a challenging task for the wide-band ICA. For this section, we applied the wide-band and the multi-band ICA on the EEG data from these participants and report the identified components time-locked to the visual stimulus.

## 4. Results

We used two variants of ICA, SOBI (second-order blind separation) [4] and InfoMax (information-maximisation) [3], to demonstrate the results in the simulations. The reason for the two choices is that, the two techniques performed differently when used for multi-band ICA, i.e., while InfoMax tends to blend PCs from different frequencies to produce the ICs, the ICs separated by SOBI are normally from a single frequency band. Other variants of the ICA had similar behavior to either of these two, for example FastICA [20] acts similar to SOBI when used for multi-band ICA. It is up to the user which flavor of the ICA to use, but we acknowledge that if the aim is to separate different sources in different frequencies, particularly the weak sources, SOBI achieves a better separation. Remixing the PCs from different frequency band is not necessarily an undesirable outcome, for example when cross-frequency coupling of the same source exists.

### 4.1. Simulation results

Figure 2 shows the identified components using wide-band and the multi-band ICA. The wide-band SOBI, Figure 2(a), separated the 5, 12, and 30 Hz sources from each other with SNRs of 11.50, 19.55, and 1.21, respectively. The IC belonging to the 30 Hz source is mixed with the lower frequency source(s). The wide-band InfoMax (Figure 2(b)) failed to separated the simulated sources from each other and the SNRs of all the identified sources was below 0.50. The multi-band SOBI, Figure 2(c), separated all three components from each other with SNRs of 11.30, 21.77, and 9.91 for the 5, 12, and 30 Hz sources, respectively. The multi-band InfoMax separated the 30 Hz source with a SNR of 9.28, but failed to separate the 5 and 12 Hz sources from each other. In fact, InfoMax mixed the PCs belonging to the 5 and 12 Hz sources which had been already separated using the the filter bank + PCA, and produced two ICs which were the result of this mixture. The topography of the two ICs also is similar to the 5 Hz source, as it is the dominant source in terms of magnitude. This behavior of InfoMax could also exist regarding the neuronal sources, that is, InfoMax could potentially mix the concurrent independent neural sources from different frequencies to create ICs. However, contrary to expectation, InfoMax did not remix the PCs belonging to the eye artifact (components 8 and 18 in Figure 2(d)) which were separated by the filter bank, to create a single component similar to IC7 in Figure 2(b). In Figure 2(b) the component IC7 is dominated by the eye artifact, although its power spectrum is not plotted, it contains small amount of power from the 30 Hz simulated source. Furthermore, IC13 contains mixture of all the simulated sources and the 50 Hz line noise, but it is dominated by lower frequency activities. In fact, the simulated 30 Hz source was scattered in several ICs when the wide-band InfoMax was used and IC13 is one of such components. Hence, removing some of these ICs such as IC7 or IC13, may result in the loss of weak neural sources. This is an example of how the wide-band component rejection could result in the loss of useful information and how multi-band signal decomposition could be a superior alternative.

Overally, multi-band SOBI performed the best in terms of separation of the simulated sources from both each other and back ground activity. Hence, we used the wide-band SOBI and multi-band SOBI for the reconstruction of real gamma-band activity in the next section.

### 4.2. Reconstruction of the gamma

For the multi-band SOBI, the filter bank was set 30-95 Hz for gamma and narrow 47-53 Hz band was defined to capture the line noise at 50 Hz. The first 45 PCs of the gamma band and the first 5 PCs of the line band were included in the global matrix of PCs (*N*_*P_gamma*_ = 45, *N*_*P_LineNoise*_ = 5). The other bands were not included in the filter bank as there the aim was to reconstruct the gamma components. As can be seen in Fig 3, narrow filter for the isolating the line noise from other components is within the band width designated for the gamma components. In this way, while we do not need to separate the gamma frequency band into below and above the 50 Hz, we can separate the line noise as a single component. If we do not allocate the narrow band, the line noise will either scatter among other components or line noise would be separated by the ICA but also includes weaker oscillations from other sources.

**Figure 2:**
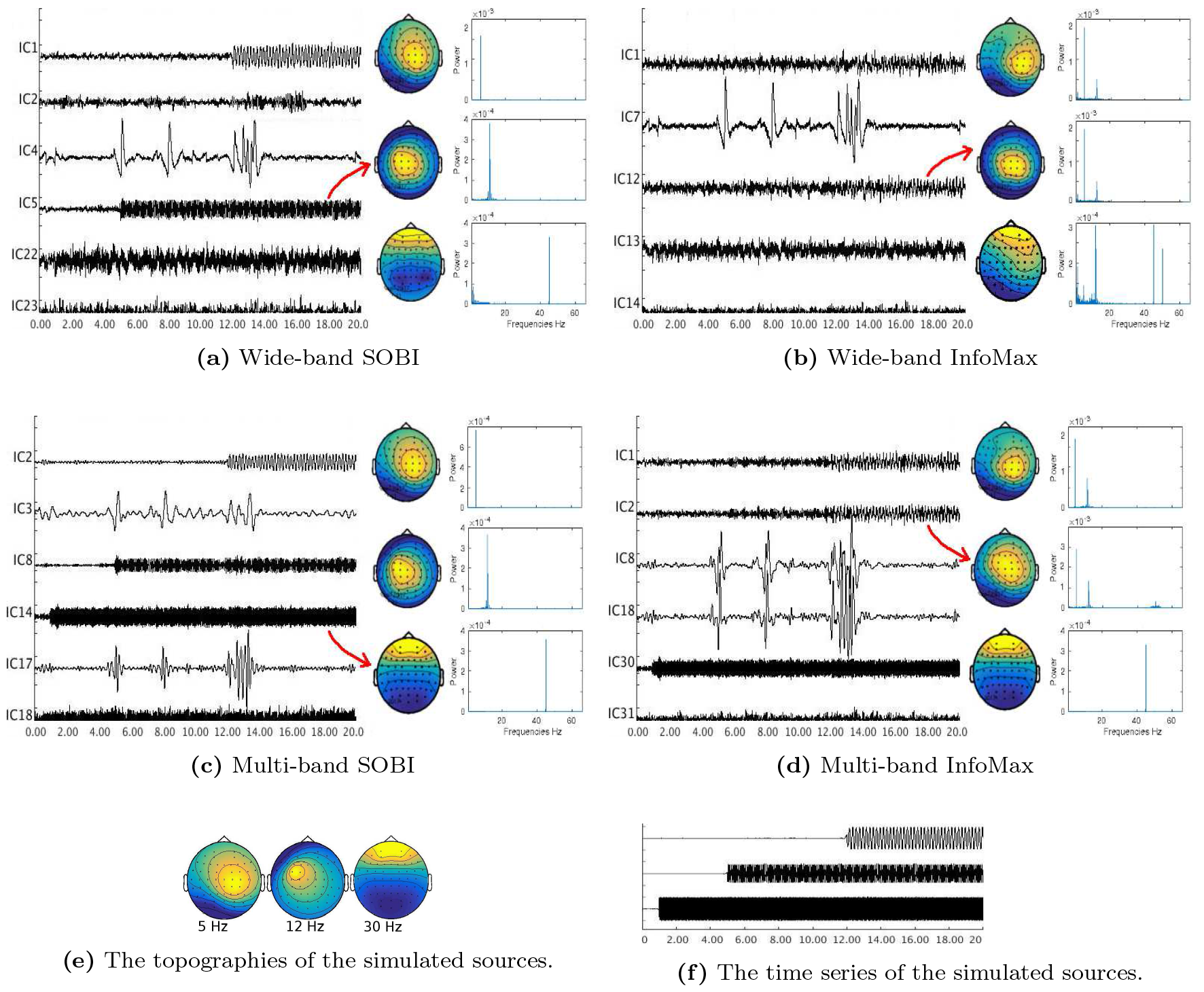
Reconstruction of the three simulated sources via wide-band SOBI (a), wide-band InfoMax (b), multi-band SOBI (c), and multi-band InfoMax (d). The groundtruth scalp maps of the 3 sources is shown in subfigure (c) and their corresponding time-courses are in subfigure (f).

Twenty out of the 22 subjects had gamma-band components time-locked to the visual stimulus with most participants having more than two components. In order to calculate the group results, the average power spectrum of the time-locked components was calculated and shown in Figure 4(a). The mean power plot of the gamma-band ICs are shown in 4(b). In order to make a mean topographic map of the all subjects and components, first the map of each IC was normalised and then its absolute value was calculated and finally summed to other maps 4(c).

**Figure 3:**
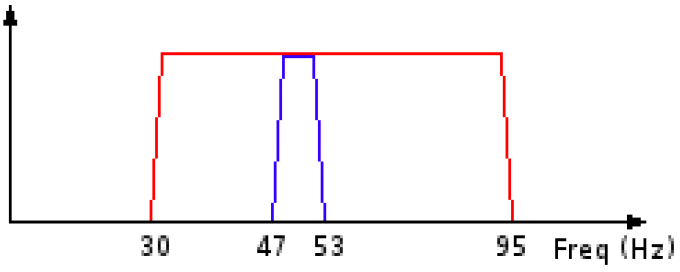
Filter bank of multi-band ICA to extract the gamma-band components.

The components separated by both the multi-band and wide-band ICA show the sustained gamma-band activation post stimulus at 0 s in Figure 4(a). However, the mean power plot of the components indicates that for wide-band ICA, the delta and alpha-band sources are the biggest contributors, whereas in multi-band ICA, the gamma-band is the primary (and only) contributor to the components. Correspondingly, the average topographic map of the components obtained by the wide-band ICA represents the dominant sources (mixture of alpha and delta-band sources) while the multi-band ICA shows the gamma-band sources. Although the maps shown in Figure 4(c) are positive only, the individual components have positive and negative polarities and Figure 4(a) shows a typical map obtained via multi-band SOBI. The posterior channels had highest contribution to the visual-induced gamma components with the right side channels having higher power. Overall, multi-band ICA allowed better reconstruction of gamma-band activity than wide-band ICA (Figure 4(a)).

## 5. Discussion

Here we introduced a novel framework, multi-band ICA, which uses a filter bank, PCA, and ICA in order to perform signal separation on the EEG (and MEG). With this approach, firstly, the wide band EEG is band-pass filtered into several subbands. Secondly, the PCA is applied on each subband to break it into spatial and temporal subspaces. Thirdly, the first few PCs of each subband which represents the strongest sources in each subband were chosen to create a global matrices of spatial and temporal PCs. Finally, ICA is applied to the global temporal PCs to estimate the temporal ICs and the mixing matrix of the ICA is then multiplied by the global spatial PCs to reconstruct the spatial ICs.

**Figure 4:**
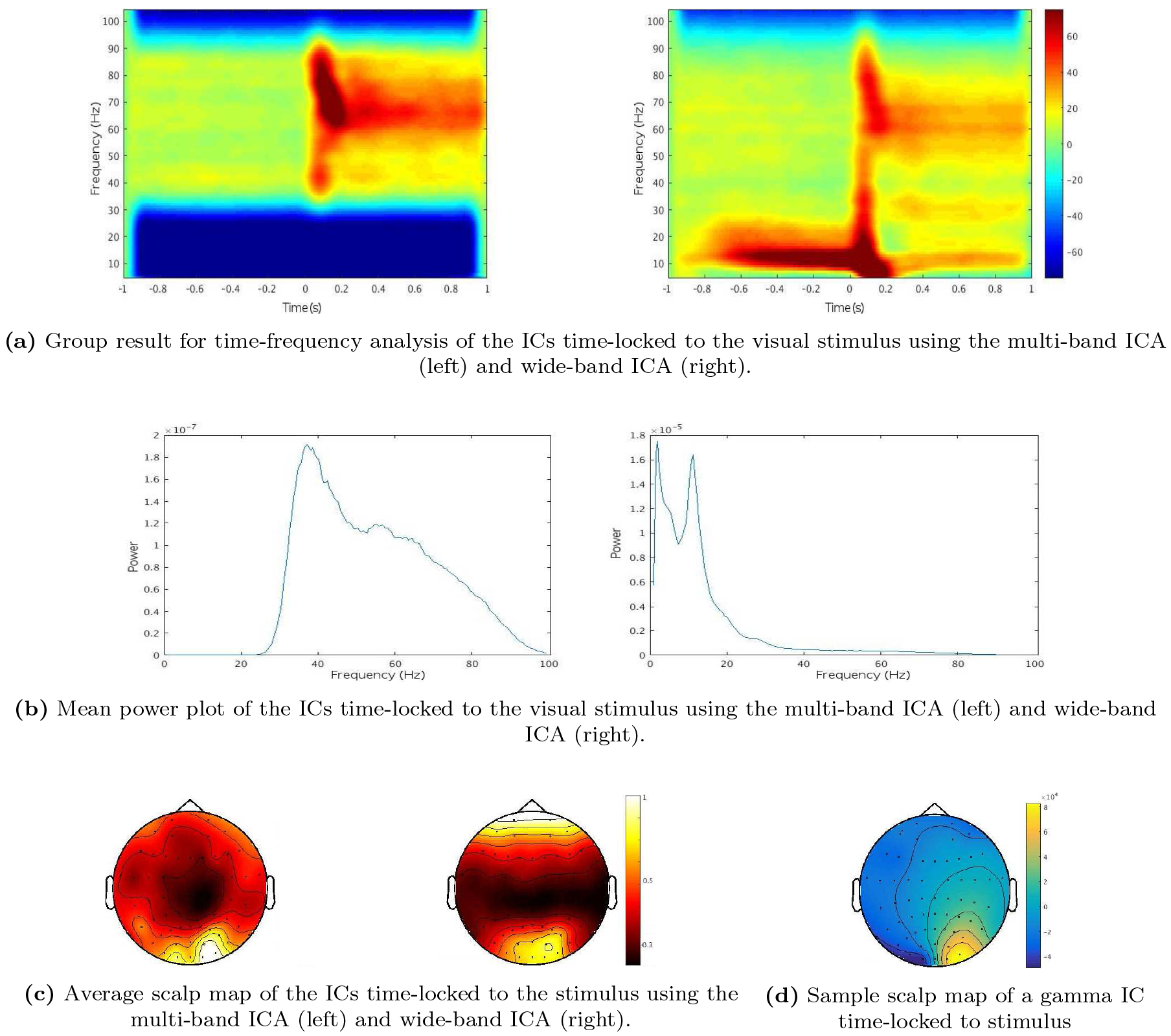
Reconstruction of the gamma-band sources time-locked to the visual stimulus using the multi-band SOBI and wide-band SOBI. The baseline for the time-frequency maps in subfigure (a) is the first 200 ms. The mean power spectrum of the components are shown in subfigure (b) and the corresponding average scalp map of the independent components are shown in subfigure (c). The map in subfigure (d) is a typical scalp map of the a time-locked gamma-band component obtained using multi-band SOBI.

Compared with the conventional wide-band ICA, the multi-band ICA is superior in terms of separation performance and the SNR of the reconstructed sources. This is due to the fact that EEG captures the activities of a large number of sources while the number of the ICs reconstructed by the wide-band ICA is limited to the number of channels. This results in the activity of the weaker sources being scattered among the ICs representing the stronger (in terms of magnitude and durations) sources. We have shown this both in the simulations and in the real gamma-band activity of subjects performing visual tasks. With the multi-band ICA, the number of ICs can be as many as number of bands × number of channels.

We did not investigate the automatic techniques such as MDL for choosing the number of the PCs from each band, and rather decided on the number of the PCs pragmatically. The automatic subspace extraction is a broad topic by its own which is beyond the scope of this paper. Based on the reconstruction of the real visual evoked sources, gamma-band components were not detected in most of the participants when we decided to extract only the first 10 PCs of the gamma-band. However, gamma-band ICs were detected in 20 out of 22 participants when we increased the number of the gamma band PCs to be included (*N*_*P_gamma*_ = 45) in the global matrix of PCs.

Our results for the study of the gamma-band sources is similar to that of [43] in which only the gamma band EEG and ICA were used for the detection of the gamma ICs. This question could be raised, rather than multi-band ICA why not band-pass the EEG and perform ICA on each band separately either for artifact rejection or for neuronal source detection. The advantage of multi-band ICA is that the user only performs visual inspection of the components once whereas the running ICA on each band separately requires corresponding visual inspections and data superimposition.

Finally, the proposed approach (multi-band ICA) can be considered as a subgroup of multi-band signal decomposition techniques, in which the aim is to reduce the decomposition burden from an adaptive technique such as ICA. The band-pass filtering of the data provides the spectral decomposition of signal mixtures and limits the task of statistical decomposition to each band. Using the concept of multi-band signal decomposition, it is possible to propose diverse subgroups both in terms of choosing a filter bank (such as wavelet filter, linear filter, non-linear filters, etc.) and a signal decomposition technique (ICA, PCA, non-negative matrix factorization, empirical mode decomposition, etc.). In this paper we used the a linear filter band + PCA + ICA to perform multi-band decompositions. Using the mentioned alternatives, optimal separations could be achieved for different types of applications.

